# A multi-seasonal model of plant-nematode interactions and its use to identify optimal plant resistance deployment strategies

**DOI:** 10.1101/774869

**Authors:** Samuel Nilusmas, Mathilde Mercat, Thomas Perrot, Caroline Djian-Caporalino, Philippe Castagnone-Sereno, Suzanne Touzeau, Vincent Calcagno, Ludovic Mailleret

**Affiliations:** Université Côte d’Azur, INRA, CNRS, ISA, France; Université Côte d’Azur, Inria, INRA, CNRS, Sorbonne Université, BIOCORE, France

**Keywords:** durable resistance, optimal strategies, resistance deployment, root-knot ne-matodes, seasonality, semi-discrete model

## Abstract

- Root-knot nematodes (RKNs) are soil-borne polyphagous pests with major impact on crop yield worlwide. Resistant crops efficiently control avirulent RKNs, but favour the emergence of virulent forms. Virulence being associated with fitness costs, susceptible crops counter-select virulent RKNs. In this study we identify optimal rotation strategies between susceptible and resistant crops to control RKNs and maximize crop yield.
- We developed an epidemiological model describing the within-season dynamics of avirulent and virulent RKNs on susceptible or resistant plant root-systems, and their between-season survival. The model was fitted to experimental data and used to predict yield-maximizing rotation strategies, with special attention to the impact of epidemic and genetic parameters.
- Crop rotations were found to be efficient under realistic parameter ranges. They were characterised by low ratios of resistant plants, and were robust to parameter uncertainty. Rotations provide significant gain over resistant-only strategies, especially under intermediate fitness costs and severe epidemic contexts.
- Switching from the current general deployment of resistant crops to custom rotation strategies could not only maintain or increase crop yield, but also preserve the few and valuable R-genes available to us.

## Introduction

As the global population increases, finding effective and durable crop protection strategies has become a major challenge (Cunniffe *et al.*, 2015). Predictions indicate that popula-growth, combined with changes in dietary habits, will lead to an increase in the global food demand by at least 50% in 2050 (Tilman *et al.*, 2011; Springmann *et al.*, 2016). To meet this demand, crop production will have to increase, with expected negative environmental impacts (biodiversity and forest loss, reduced freshwater availability, soil degradation and CO2 emissions) if relying on the extensive use of chemical pesticides and monocultures (Tilman *et al.*, 2001; Stoate *et al.*, 2009; Zhan *et al.*, 2015). Furthermore, crop losses are expected to increase as well, owing to the emergence or evolution of plant pests and diseases (Palumbi, 2001; Stukenbrock & McDonald, 2008). These trends call for experimental and theoretical studies aiming at protecting crops and increasing their yield durably, while reducing pesticide dependence. In this context, the development of environmentally-friendly pest management strategies based on biological control, better cultural practices and the use of resistant plants are very promising (Mundt, 2014; Zhan *et al.*, 2015; Van Lenteren *et al.*, 2018).

Natural plant resistance is amongst the most efficient alternatives to pesticides in economic, environmental and social terms (Biffen, 1905; Hammond-Kosack & Jones, 1997). Qualitative plant resistance rests on gene-for-gene interactions (Flor, 1971), in which an avirulent gene (Avr-gene) in the pest or pathogen interacts with a major resistance gene (R-gene) in the plant, resulting in disease resistance through what is usually called effector-triggered immunity or incompatible reaction (Dangl & Jones, 2001; Jones & Dangl, 2006). If the R-gene is inactive or absent, or equivalently if the pest lacks the Avr-locus, the interaction instead results in plant infection. Major R-genes are rare in nature and plant breeders mostly work on the introgression of a small list of major R-genes into different genetic backgrounds to create commercial crop cultivars. Therefore, farmers ultimately employ the same resistance genes over several years and on large spatial scales. Such an intensive use of resistance generates strong selection pressures on populations of avirulent pests, that can lose the Avr-gene through mutation, causing the emergence and establishment of virulent variants (Leonard, 1977; Castagnone-Sereno, 2002; McDonald & Linde, 2002; Parlevliet, 2002; Garcia-Arenal *et al.*, 2003).

According to Johnson (1981), a durable resistance is one that remains effective in a cultivar for a long period of time despite its widespread cultivation. Resistance durability may depend on the time required for a mutation at the Avr-gene to occur and the time for the virulent pathogen to establish (Van den Bosch & Gilligan, 2003; Stuthman *et al.*, 2007; Barrett *et al.*, 2008; Fabre *et al.*, 2009; Brown, 2015; Zhan *et al.*, 2015). The latter might be expected to be very short, considering the huge advantage for a pathogen to overcome resistance and become virulent. However, significant polymorphism exists at virulence genes, that can at least partly be explained by fitness costs associated with virulence (Stahl *et al.*, 1999; Tian *et al.*, 2003; Laine & Tellier, 2008). Numerous studies have reported fitness costs in bacteria (Cruz *et al.*, 2000; Leach *et al.*, 2001), oomycetes (Montarry *et al.*, 2010) or viruses (García-Arenal & Fraile, 2013). The existence of fitness costs implies that even though virulent pathogens are selected for in resistant crops, they are selected against in susceptible crops, where avirulent pathogens grow and reproduce faster.

Several approaches to improve the durability of R-genes that are based on suitable combinations of resistant and susceptible plant cultivars have been proposed (Van den Bosch & Gilligan, 2003; Fabre *et al.*, 2012; Papaïx *et al.*, 2014; Fabre *et al.*, 2015; Lof & van der Werf, 2017). The most common deployment strategies are mixtures, mosaics and rotations, that exploit spatial and/or temporal heterogeneity in selection pressures (Kiyosawa, 1982; Mundt, 2002; Pink, 2002; Djidjou-Demasse *et al.*, 2017; Rimbaud *et al.*, 2018). Crop rotation is one of the most important strategies thought to delay and control resistance breakdown when a single R-gene is deployed over a large area (Dury *et al.*, 2012). Root-knot nematodes (RKNs, *Meloidogyne spp.*) are ubiquitous plant pathogens (Trudgill & Blok, 2001; Jones *et al.*, 2011). They are obligate extremely polyphagous plant endoparasites, that cause damages to the roots of thousands of host plant species (Wesemael *et al.*, 2011; Perry *et al.*, 2010). Overall, their economic impact has been estimated at over 121 billion dollars of crop losses each year (Chitwood, 2003). For several decades, controlling these parasites has relied on chemical treatments, but these proved extremely damaging to the environment and to human health and have been banned (Zasada *et al.*, 2010; Abad & Williamson, 2010). The fight against RKNs is now largely based on the use of plant cultivars bearing resistance genes (Williamson & Roberts, 2010). However, resistance breaking by virulent nematodes has been demonstrated in the laboratory (Jarquin-Barberena *et al.*, 1991; Djian-Caporalino *et al.*, 2011; Meher *et al.*, 2009) and is more and more observed in field conditions (Verdejo-Lucas *et al.*, 2009). As for other plant parasites, virulence in RKNs is associated with a fitness cost, and it was shown that virulence reduces the capacity to infect the plant, as well as the number of eggs laid per female (Castagnone-Sereno *et al.*, 2007; Djian-Caporalino *et al.*, 2011). Therefore, setting up rotation strategies of resistant and susceptible cultivars has the potential to increase the durability of resistance genes and the efficacy of resistance–based nematode control. However, field tests of deployment strategies in terms of epidemiological control and resistance durability remain difficult, owing to their labor intensive nature and to the long time horizons involved (Djian-Caporalino *et al.*, 2014).

In these conditions, modelling approaches constitute a powerful way to explore resistant plant deployment strategies and assess their efficiency to reduce yield losses and increase control durability (Brown, 2015; Papaïx *et al.*, 2018). Unfortunately, the literature is very poor in theoretical modelling studies addressing the control of soil-borne pathogens with limited dispersal, such as RKNs. For instance, most studies deal with pathogens that can disperse over large spatial scales (Gilligan, 1995; Thrall *et al.*, 1997; Otten & Gilligan, 2006; Fabre *et al.*, 2012; Djidjou-Demasse *et al.*, 2017; Lof & van der Werf, 2017). RKNs, in contrast, have very limited mobility in the soil, feeding and reproducing locally in the plant root system. Consequently, nematode populations barely mix and strategies based on spatial arrangements are poorly applicable. In addition, the major RKN species reproduce solely by clonal reproduction so that techniques based on recombination between virulent and avirulent genotypes do not operate. The purpose of this study was to assess quantitatively whether rotation strategies between susceptible and RKN-resistant cultivars can be efficient to control RKNs, and to determine which optimal crop rotation strategies should be used to maximise crop yield over several seasons. We did this by building a semi-discrete plant epidemic model (Fabre *et al.*, 2012; Mailleret & Lemesle, 2009; Mailleret *et al.*, 2012), tailored to the RKN pathosystem. The model describes the within-season dynamics of the interaction between a plant root system and RKNs, the owerwintering dynamics between consecutive seasons and the potential evolution of the nematode population from avirulent to virulent forms. The model was parameterized from the literature and fitted to experimental data (Ehwaeti *et al.*, 1998). We used the model to compute optimal crop rotation strategies with respect to a proxy of crop yield over different time horizons. Given that the fitness costs vary among R-genes and nematode strains, and are crucial to the durability of R-genes, we payed special attention to the influence of these genetic parameters. We evaluated to what extent crop rotation provided better crop yield than the widely used resistant plant-only strategy (pure resistant strategy) for different epidemiological scenarios and genetic parameters. We also tested the robustness of our results to determine whether the effectiveness of optimal periodic rotations can be maintained even if epidemiological and genetic parameters are not known precisely. We investigated the key factors to be taken into account for optimal resistance plant deployment strategies against RKNs. Our results showed that crop rotation can very often increase crop yield significantly, in all scenarios tested. The efficacy of optimal rotation strategies also proved robust to parameter uncertainty; specifically, optimal rotations consistently outperformed the resistant-only strategy.

## Model description

### Study system

We focused on root-knot nematodes (RKNs) of the species *Meloidogyne incognita*. These are obligate endoparasites of plant roots, and reproduce only by clonal reproduction. *M. incognita* is one of the most prevalent species in the warm conditions of Mediterranean countries, especially in protected crops (Wesemael *et al.*, 2011). The life cycle of *M. incognita* consists in four stages that can be achieved in three to five weeks, depending on environmental conditions (Abad & Williamson, 2010). Second-stage juveniles dwell in the soil and penetrate the plant when a root grows in their vicinity. Once a nematode reaches the vascular cylinder of the root, salivary secretions induce the creation of a feeding site. These are composed of five to six hypertrophied plant cells, known as giant cells. The nematode spends the rest of its life in this feeding site, where it develops until reproduction. When mature, adult females release several hundreds of eggs (between 300 to 2000 eggs/female on average) outside the root, that will hatch into free living juveniles and complete the cycle (Fig. 1).

**Fig. 1.**
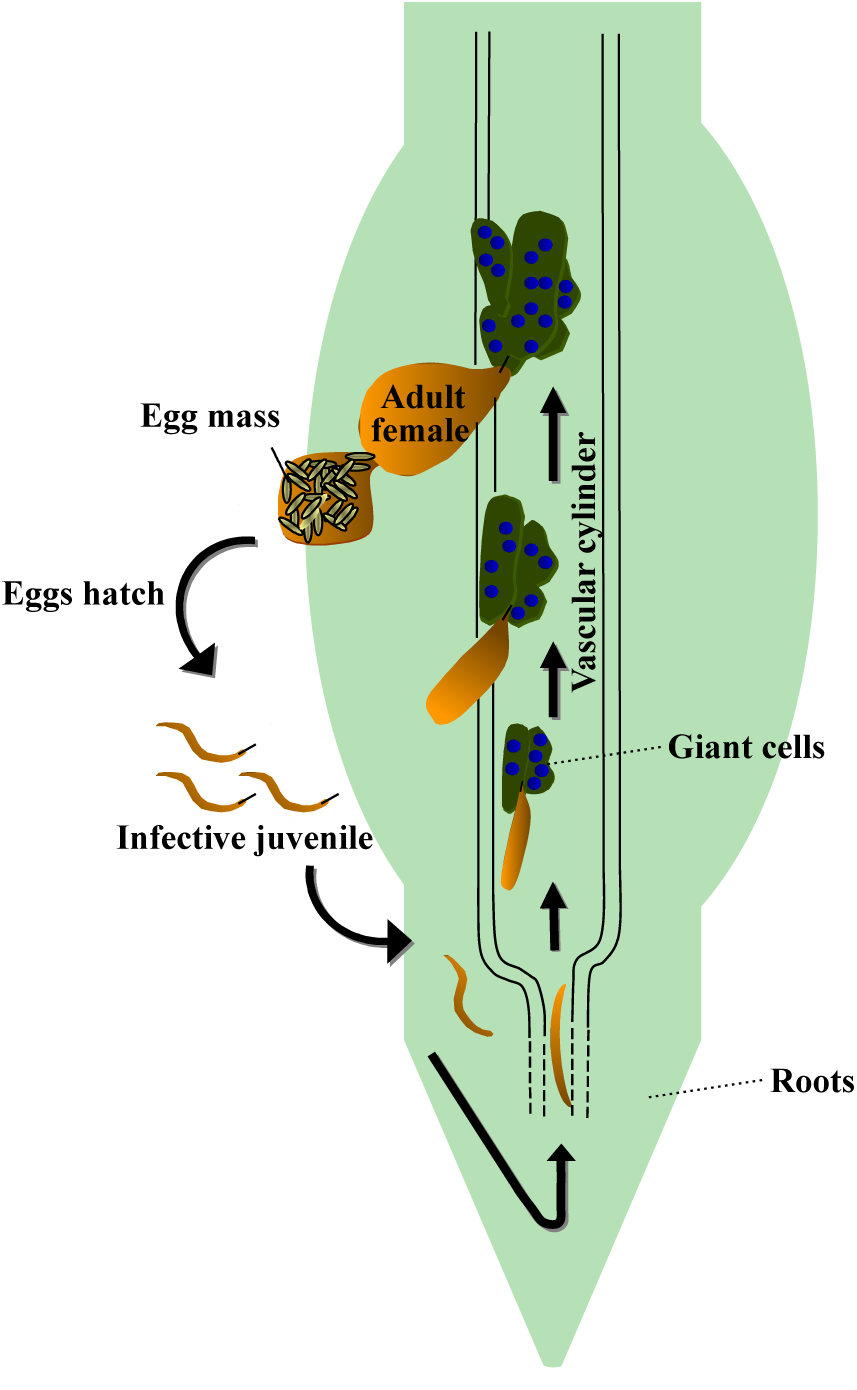
The life cycle of root-knot nematodes – adapted from Williamson & Gleason (2003) and Abad & Williamson (2010).

In the *Solanaceae* plant family, a few resistance genes are known to block the development and reproduction of RKNs: the *Mi-1* gene in tomato (Milligan *et al.*, 1998) and the *N, Me-1* and *Me-3* genes in sweet pepper (Djian-Caporalino *et al.*, 2007, 2011). The most pervasive resistance breakdown issue consists in the *Mi-1* gene being overcome by *M. incognita* (Ornat *et al.*, 2001; Seid *et al.*, 2015). *Mi-1*, originally from the wild species *Solanum peruvianum*, was introgressed into tomato by interspecific crosses in the early 1940s. The first resistant varieties appeared on the market by the end of that decade. Since then, many resistant varieties have been globally deployed, all bearing the same resistance gene. Nowadays, resistance breaking by *M. incognita* populations is recorded worldwide, in virtually every area growing tomatoes (Seid *et al.*, 2015). In this study, we will thus use *Mi-1*: *M. incognita* as our reference system.

### Model of plant-nematode interactions

The interaction between nematodes and plants during a cropping season was modeled as an epidemic of free living pests infesting and spreading among the plant root system. We first consider only avirulent nematodes and a susceptible plant. The model describes in continuous time the changes in four variables: the density of free living nematodes in the soil (*P*_*a*_), the density of healthy susceptible plant roots (*H*^*S*^) and the density of latent (*E*_*a*_) and infectious (*I*_*a*_) feeding sites induced by nematodes. It is represented by the following system of differential equations:

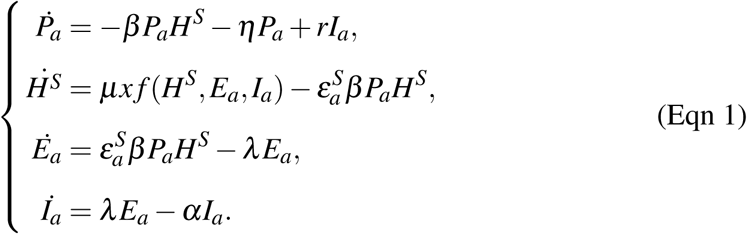

When a free living avirulent nematode *P*_*a*_ comes into contact with a portion of healthy plant root *H*^*S*^, the latter becomes latently infected *E*_*a*_ at rate 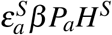, where *β* is the infection rate and 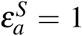 is a conversion factor between nematode and root densities (Table 1). After a time period 1*/λ*, the infected root portion becomes infectious (*I*_*a*_) and starts producing free living avirulent nematodes (*P*_*a*_) at rate *r*. Free living nematodes in the soil and infectious nematodes in the roots die at rates *η* and *α*, respectively. Roots are assumed to grow linearly with time at basic rate *µx* (Leskovar *et al.*, 1990), where *x* is a conversion factor between root biomass and root density. Root infection by nematodes is known to impact root growth (Zeck, 1971), which is taken into account through function *f* (.). This function discounts the basic growth rate by a decreasing exponential function of infection prevalence 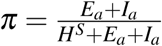 multiplied by a scaling factor *k*: *f* (*H*^*S*^, *E*_*a*_, *I*_*a*_) = *e*^−*kπ*^. The model (Eqn 1) is readily extended to take into account susceptible and resistant plants, as well as the co-occurence of avirulent and virulent nematodes. Variable *P*_*v*_ represents the density of virulent free living nematodes in the soil; similarly *I*_*v*_ and *E*_*v*_ represent the densities of feeding sites infected by latent and infectious virulent nematodes, respectively. In what follows, superscript *X* indicates the type of cultivated plant in the current cropping season, *i.e.* either susceptible (*X* = *S*) or resistant (*X* = *R*). The model then reads:

**Table 1.** Model variables and parameters.

| Symbol | Description | Value(s) | Unit | Ref. |
| --- | --- | --- | --- | --- |
| $H^X$ | Density of healthy plant root | | UR | |
| $P$ | Density of free living nematodes | | UN | |
| $E$ | Density of latently infected feeding sites | | UR | |
| $I$ | Density of infectious feeding sites | | UR | |
| $H_0$ | Initial root biomass | 6 | UR | [1] |
| $P_0$ | Initial nematode density in the soil | 0.8 (0.03, 20) | UN | [2] |
| $p_v$ | Initial proportion of virulent nematodes | $10^{-3}$ ($7 \cdot 10^{-4}$, $1.3 \cdot 10^{-3}$) | – | [3] |
| $\beta$ | Infection rate | $1.11 \cdot 10^{-4}$ ($7.78 \cdot 10^{-5}$, $1.44 \cdot 10^{-4}$) | $\text{UR}^{-1} \text{ day}^{-1}$ | [*] |
| $w_\beta$ | Fitness cost on infectiveness | 0.09 (0.06, 0.12) | – | [4] |
| $\lambda$ | Transition rate from $E$ to $I$ | 0.06 | $\text{day}^{-1}$ | [4,5] |
| $r$ | Nematode reproduction rate | 17 (11.9, 22.1) | $\text{UN UR}^{-1} \text{ day}^{-1}$ | [4] |
| $w_r$ | Fitness cost on reproduction | 0.31 (0.22, 0.40) | – | [4] |
| $\delta$ | Fraction of virulent offspring | $10^{-6}$ ($7 \cdot 10^{-7}$, $1.3 \cdot 10^{-6}$) | – | [3] |
| $\alpha$ | Nematode mortality rate in roots | 0.125 (0.0875, 0.1625) | $\text{day}^{-1}$ | [4,5] |
| $\eta$ | Nematode mortality rate in the soil | 0.04 (0.028, 0.052) | $\text{day}^{-1}$ | [6] |
| $\phi$ | Between-season survival probability | 0.4 (0.28, 0.52) | – | [7] |
| $\varepsilon_y^X$ | Nematode infection success | 0 if $X = R$ and $y = a$<br>1 otherwise | $\text{UR UN}^{-1}$ | |
| $\mu$ | Plant root growth rate | 105 | $\text{mg day}^{-1}$ | [1] |
| $x$ | Conversion factor between root mass and density of feeding sites | $3 \cdot 10^{-3}$ | $\text{UR mg}^{-1}$ | [*] |
| $k$ | Impact of infection prevalence on root growth rate | 10.33 | – | [*] |
| $\tau$ | Duration of a cropping season | 135 | days | [8] |

**Bold parameters** were varied for the sensitivity analysis: default value and $\pm 30\%$ variations (indicated in brackets) were tested; larger variations were tested for $P_0$ , in line with Ehwaeti *et al.* (1998).
**Units:** UR: number of feeding sites per gram of soil; UN: number of nematodes per gram of soil.
**Sources:** [1] Leskovar *et al.* (1990); [2] Ehwaeti *et al.* (1998); [3] Ploeg & Maris (1999); [4] Castagnone-Sereno *et al.* (2007); [5] Ekanayake (1986); [6] Tsai (2008); [7] Castagnone, unpublished data; [8] Djian-Caporalino, unpublished data; [8] Djian-Caporalino, unpublished data; [\*] Estimated.

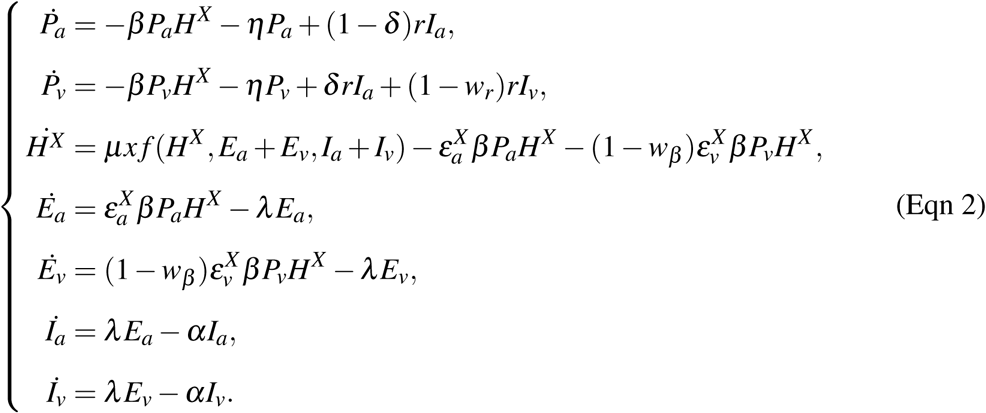

Avirulent and virulent nematodes compete for healthy plant roots *H*^*X*^ in the following way: avirulent nematodes *P*_*a*_ can infect susceptible plants 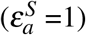 but are unable to infect resistant plants 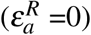, while virulent nematodes *P*_*v*_ are able to infect both resistant 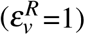 and susceptible plants 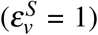 Importantly, virulent nematodes grow more slowly than avirulent ones because they suffer from fitness costs, at two levels: reduced infectiveness (*w*_*β*_) and reduced reproduction (*w*_*r*_) (Jarquin-Barberena *et al.*, 1991; Castagnone-Sereno *et al.*, 2007; Meher *et al.*, 2009; Djian-Caporalino *et al.*, 2011). We considered that there was no additional fitness cost (also called “residual effect”) on resistant plants. Indeed, we conducted statistical tests and found no significant differences in terms of fitness costs when virulent nematodes grew on resistant *Mi-1* or susceptible tomato plants (Castagnone-Sereno *et al.*, 2007). Furthermore, we assumed that a fraction *δ* of avirulent nematode offspring are virulent (Castagnone-Sereno *et al.*, 1994), due to mutation and/or epigenetic mechanisms. We also postulated that, once acquired, virulence could not be lost by the virulent lineage (Castagnone-Sereno *et al.*, 1993). We can thus charaterize a resistance gene and its suceptibility to resistance breakdowns with a set of three genetic parameters that: the two fitness costs associated with nematode virulence (*w*_*β*_ and *w*_*r*_) and the proportion of virulent variants in the nematode offspring (*δ*).

The initial conditions of the full multi-seasonal model were set to *H*^*X*^ (0) = *H*_0_, the initial root biomass of newly planted individuals, *P*_*a*_ = (1 − *p*_*v*_)*P*_0_ and *P*_*v*_ = *p*_*v*_*P*_0_, where *P*_0_ refers to the initial nematode density in the soil and *p*_*v*_ to the initial proportion of virulent nematodes in the soil. Initial values of *I*_*a*_, *E*_*a*_, *I*_*v*_ and *E*_*v*_ were set to 0 because plants were assumed to be healthy at the time they were planted.

At the end of each cropping season, plants are removed. At the beginning of the next cropping season, healthy and infected roots are thus reset to their initial values, *H*_0_ and 0, respectively. Nematode densities *P*_*a*_ and *P*_*v*_ are set to their value at the end of the previous cropping season, multiplied by a survival probability *ϕ*. The full model of plant nematode interaction over multiple cropping seasons is therefore a hybrid model, with a continuous part to describe the nematode infection dynamics during a cropping season of length *τ*, and a discrete part to describe nematode survival between seasons and crop planting (Mailleret & Lemesle, 2009; Mailleret *et al.*, 2012). The model is represented graphically in Supporting Information Fig. S1.

For simulations and numerical investigations, models (Eqn 1) and (Eqn 2) were implemented using the R software (http://www.r-project.org) and ordinary differential equations were solved with the DESOLVE R package (https://CRAN.R-project.org/package=desolve). We also analyzed the existence and stability of the nematode-free stationary solution and computed the season to season basic reproduction numbers *R*_0_ for avirulent and virulent nematodes (Mailleret *et al.*, 2012). *R*_0_ computations are detailed in Supporting Information Methods S1.

### Model parameterization and epidemiological scenarios

Most parameter values could be set from published estimates in the literature (Table 1). No data were available for three parameters: the infection rate (*β*), the conversion factor between root biomass and density of feeding sites (*x*) and the plant growth scaling factor (*k*). Their values were thus estimated by fitting model (Eqn 1) to an experimental dataset reporting the final nematode density in plant roots as a function of initial nematode density in the soil (Ehwaeti *et al.*, 1998). Specifically, avirulent *M. incognita* nematodes were inoculated at controlled densities in the soil, then tomato plants (cv *Moneymarker*) were planted and the nematode density in the root system was measured after 42 days and 135 days of cultivation. The relative root biomass (*i.e.* root biomass divided by the control root biomass with no nematode) was also measured. Both measurements after 135 days were used to fit our model, and the measurements at day 42 were compared to predicted values to assess model validity (Fig. 2). More details are available in Supporting Information Methods S2.

**Fig. 2.**
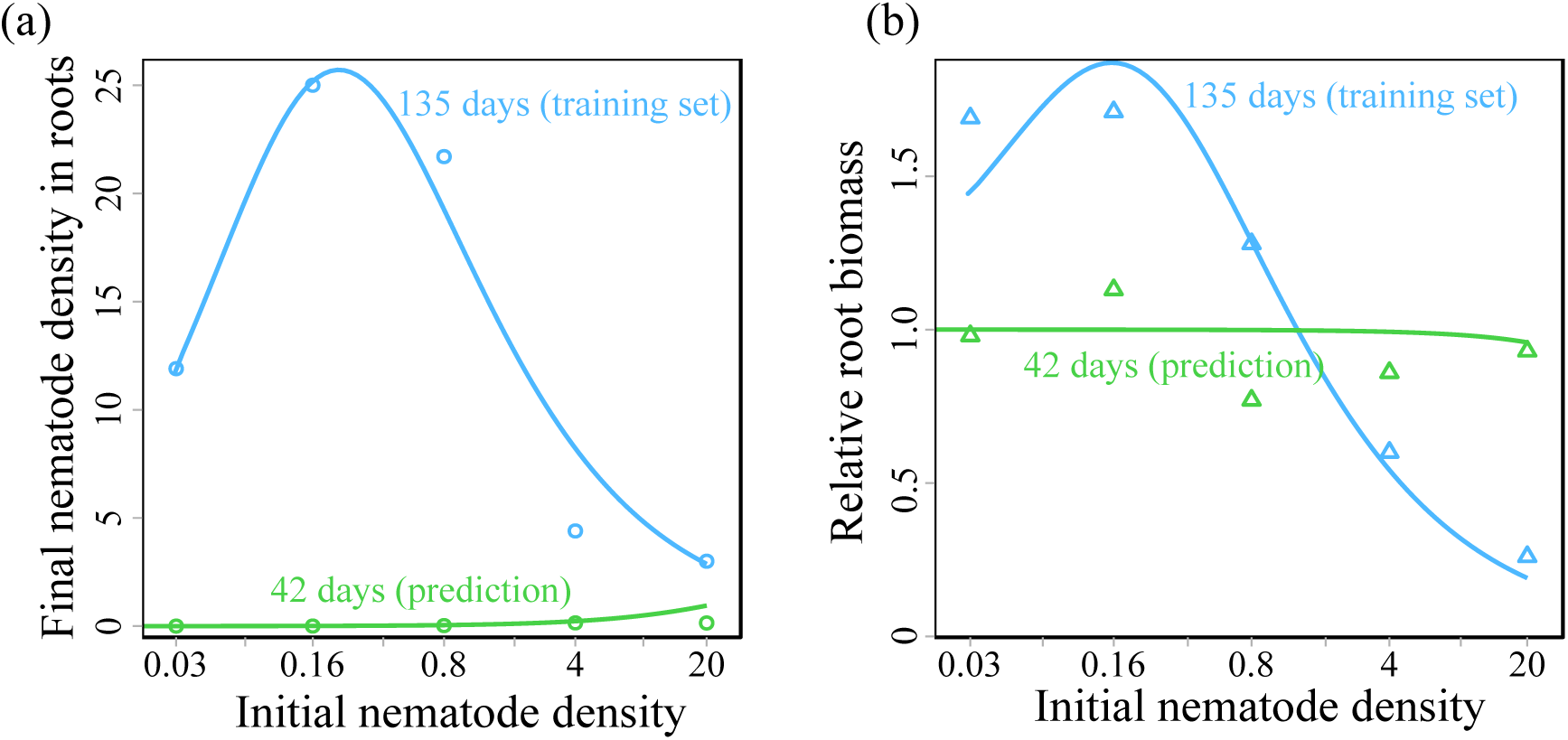
Fit of the model to experimental data over one cropping season (Ehwaeti *et al.*, 1998). **(a)** Final density of nematodes in the roots and **(b)** relative biomass after 42 (in green) and 135 days (in blue), as functions of the initial density of nematodes in the soil (log scale). Model outputs are shown as solid curves, circles and triangles represent experimental measurements.

Parameters characterising virulent nematodes, *i.e.* the fitness costs, were selected from data on the *Mi-1* resistant tomato Castagnone-Sereno *et al.* (2007). All parameters are summarised in Table 1.

From the estimated parameter values, we performed a global sensitivity analysis (Saltelli *et al.*, 2008) and defined four contrasted epidemiological scenarios. We carried out the analysis with the multi-seasonal model (Eqn 2), over 15 seasons of susceptible plant cultivation. We varied epidemiological parameters (values given in Table 1) and determined their impact on a crop yield proxy, the healthy root density 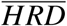 defined below in (Eqn 3). More details are available in Supporting Information Methods S3. The most influential parameters were found to be the nematode reproduction rate *r*, the nematode mortality rate in the soil *η*, the infection rate *β* and the nematode mortality rate in the root *α*. By varying these parameters around their estimated values, we created four epidemiological scenarios, corresponding to different levels of nematode agressiveness, form low to extreme (Table 2).

**Table 2.**
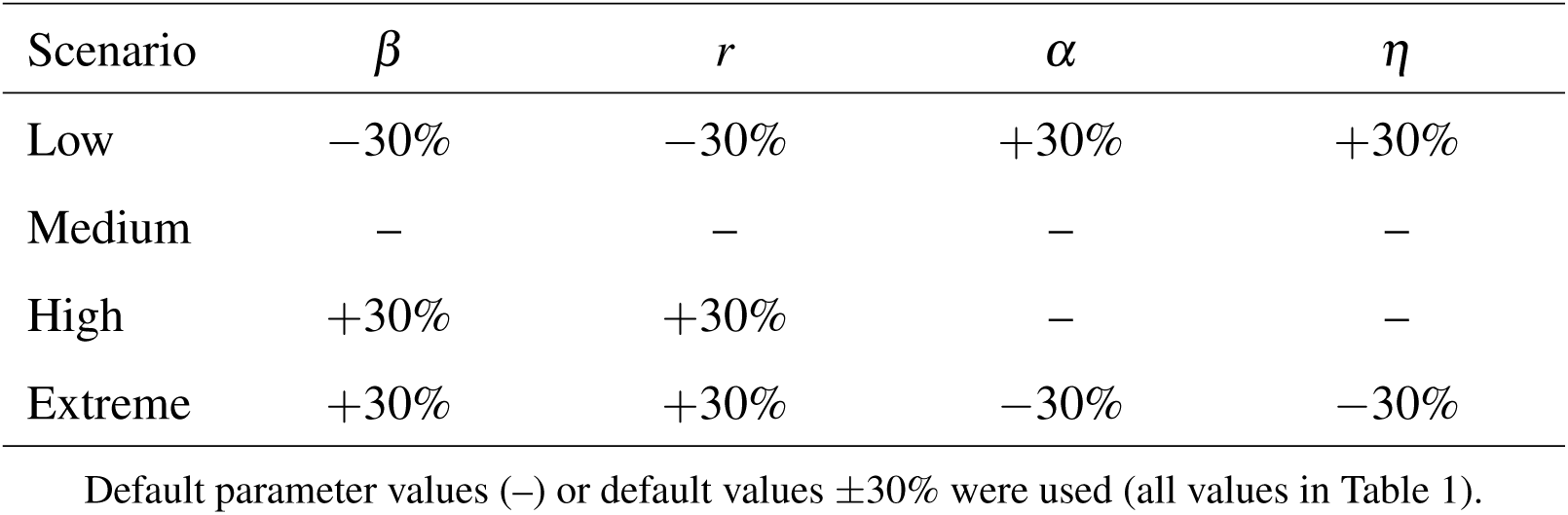
Definition of the four epidemiological scenarios based on the four most influential epidemiological parameters: nematode infection rate (*β*), reproduction rate (*r*), mortality in the roots (*α*) and in the soil (*η*).

### Resistance deployment strategies

We considered several resistance deployment strategies: the two “pure” resistant-only and susceptible-only strategies, consisting in planting one crop type all the time; periodic rotation strategies, alternating resistant and susceptible plants according to a repeated pattern; and unconstrained strategies, *i.e.* arbitrary sequences of susceptible and resistant plants.

The performance of each strategy was quantified with the “healthy root density” 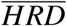, a proxy of crop yield defined as the mean of the integral of healthy plant root densities over the *n* cropping seasons:

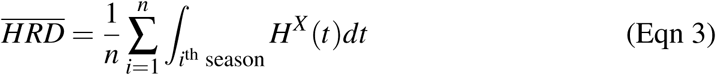

This quantity is similar to the healthy leaf area duration (*HAD*), the integral of healthy green canopy area during a growing season, used by many authors for airborne pathogens (Waggoner & Berger, 1987; Gooding *et al.*, 2000; Van den Bosch & Gilligan, 2003; Lo Iacono *et al.*, 2012; Elderfield *et al.*, 2018; Papaïx *et al.*, 2018).

The optimal strategy over a given time horizon of *n* cropping seasons was defined as the strategy with best performance, *i.e.* maximising the 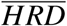 (Eqn 3). To identify optimal periodic strategies, we computed all periodic rotation strategies between resistant and susceptible crops, beginning with resistant crops and alternating *m* and *p* seasons of resistant and susceptible plants, respectively. We denoted by *mR* + *pS* these periodic strategies. As an example, Fig. 3a displays the 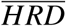 of all periodic rotation strategies over a 15-season time horizon. The optimal periodic strategy is 1*R* + 5*S*, which corresponds to 1 season of resistant plants followed by 5 seasons of susceptible plants, and so on. A graphical representation in Fig. 3b–d displays the nematode and plant root dynamics over time. We also identified unconstrained optimal strategies by using a genetic algorithm implemented through the GENALG R package (https://CRAN.R-project.org/package=genalg). We computed these optimal strategies and the corresponding ratios of resistant plants, *i.e.* the number of seasons with resistant plants divided by the given time horizon, for all time horizons between 1 and 30 cropping seasons.

**Fig. 3.**
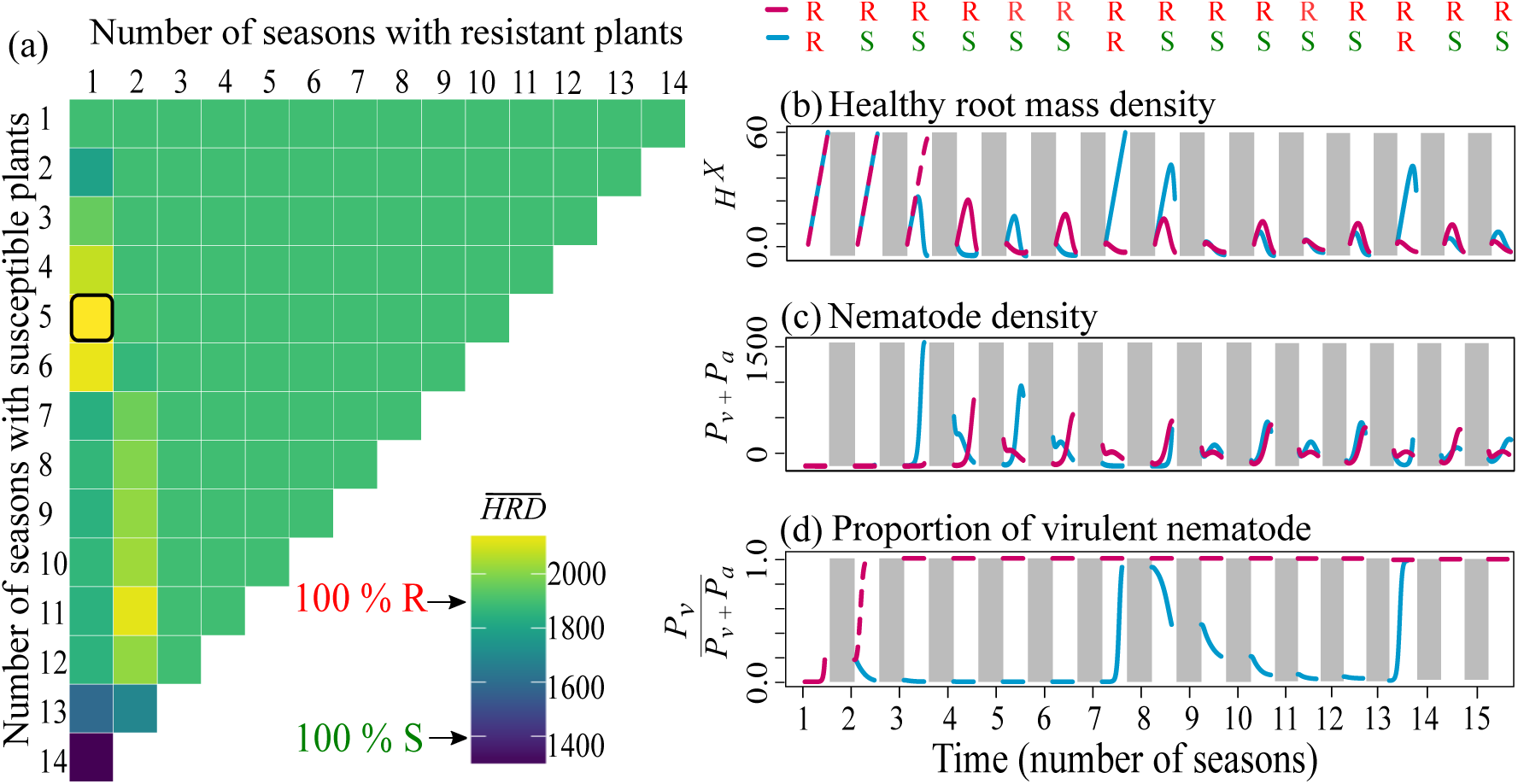
**(a)** Performance (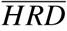, colour scale) of all periodic rotation strategies, according to their number of seasons of resistant (in columns) and susceptible (in rows) plants, over a 15-season time horizon; performance of the susceptible-only and resistant-only strategies are indicated on the color scale; the black-framed strategy is the optimal strategy 1*R* + 5*S*. **(b–d)** Graphical representation of two strategies: the resistant-only strategy (in pink) and the 1*R* + 5*S* periodic strategy (in blue), which is optimal over a 15-season time horizon; shaded areas correspond to the inbetween seasons. Default parameter values were used (Table 1).

We analysed with particular attention the influence of the genetic parameters (fitness costs *w*_*β*_, *w*_*r*_ and the proportion of virulent offspring *δ*), possibly in combination with the epidemiological scenarios, on the nature and relative performance of optimal rotation strategies. Specifically, we sought to determine when optimal rotation strategies could outperform the usual resistant-only strategy, and to what extent crop yield could be increased by using such rotation strategies.

Finally, we evaluated the robustness of our results to determine to what extent optimal periodic strategies would remain effective if biological parameters were not known with perfect precision. For the medium, high and extreme epidemiological scenarios defined in Table 2, the optimal periodic strategy was computed over a 15-season time horizon and its performance was tested against *±*10% variations in the values of six key parameters (*ϕ, w*_*r*_, *w*_*β*_, *P*_0_, *p*_*v*_ and *δ*), on top of the four epidemiological parameters defining the scenarios. In contrast with the analysis focusing on the impact of the genetic parameters, the periodic strategy was not computed afresh when the key parameters varied. For each epidemiological scenario, we explored this parameter space using a fractional factorial design containing 243 parameter combinations. The design was obtained using the PLANOR R package (https://CRAN.R-project.org/package=planor).

## Results

### Optimal deployment strategies

The performance 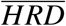 of the four types of deployment strategies are shown in Fig. 4a, for different time horizons and the default parameters. As expected, the resistant-only and optimal strategies outperformed the susceptible-only strategy, since the deployment of resistance prevents infection by avirulent nematodes. However, for these strategies, the 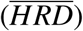 decreased with the time horizon. This is also expected, as the deployment of resistance also causes virulent nematodes to appear and take over the nematode population.

**Fig. 4.**
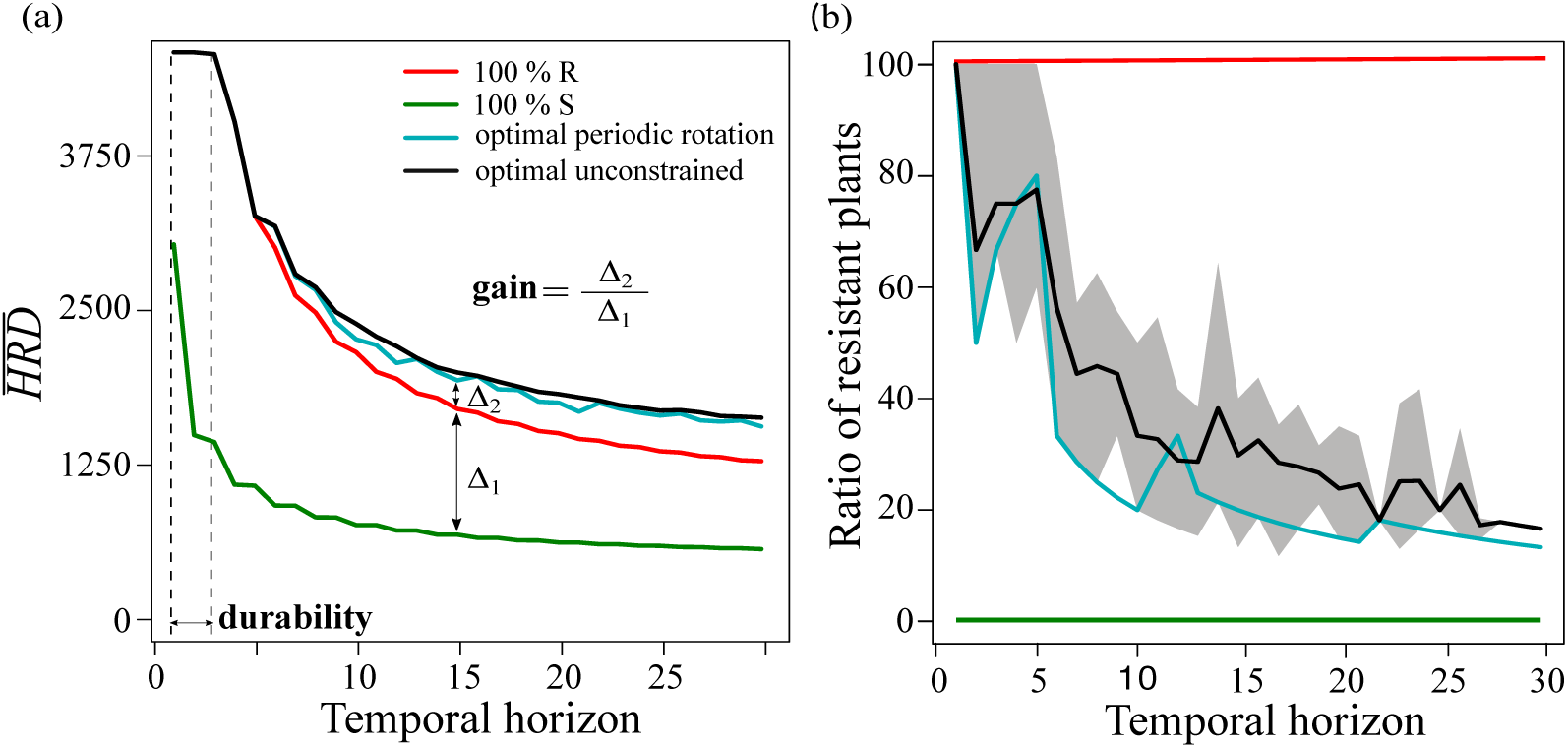
**(a)** Healthy root density 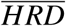 and **(b)** ratio of resistant plants as functions of the time horizon, for different deployment strategies: susceptible-only (green), resistant-only (red), optimal periodic rotation (blue) and optimal unconstrained (black). Different unconstrained optimal strategies (yielding the same 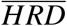) were identified, so the ratio of resistant plants is represented in panel (b) by its range (shaded area) and its average value (black solid curve). Default parameter values were used (Table 1). The durability (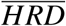 loss *<* 1%) and the relative gain, defined more thoroughly in the text, are illustrated in panel (a).

For up to five years of cultivation, the resistant-only strategy performed as well as any optimal deployment strategy, but over longer time horizons, it could be significantly outperformed. For instance, over 15 cropping seasons the 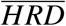 was around 2044 with optimal strategies, while it had dropped to 1822 for a pure R strategy (Fig. 4a). We also remark that, for all time horizons considered (up to 30 years), the optimal periodic and the unconstrained strategies had almost identical performances. This indicates that periodic rotations are almost optimal in this system.

The deployment of a pure resistant-only strategy is thus reasonable for at most five years in this cropping system. Beyond that, the optimal strategy generally was to alternate one season of resistant plants with a few seasons of susceptible plants, as shown for instance in Fig. 3 for a 15-season time horizon (Fig. 4b). This optimal strategy ensures that virulent nematodes remain sufficiently rare in the soil, allowing to sustain the efficiency of resistant plants, which severly reduce the avirulent nematode population. The average ratio over the time horizon for the unconstrained optimal strategy was generally more important than for the optimal periodic rotation strategy (Fig. 4b). For instance, over a 15-season time horizon, the genetic algorithm identified 11 equivalent solutions and the ratio of resistant plants deployed was on average 30%. For the optimal periodic strategy, it was only 25%. Unconstrained optimal strategies identified by the genetic algorithm are actually fairly similar to optimal periodic rotations in terms of structure, except that more resistant plants are used in the final seasons, explaining the higher ratio of resistant plants in unconstrained strategies.

Fig. 4a illustrates the definition of two synthetic metrics that are used in this paper. The first one is the relative gain, defined as the 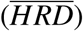 gain obtained by shifting from a resistant-only to an optimal strategy, relative to the gain obtained by shifting from a susceptible-only to a resistant-only strategy. For a given number of cropping seasons (*i.e.* a given time horizon), a positive relative gain indicates that the optimal periodic rotation strategy outperforms the resistant-only strategy, whereas a negative value indicates that the resistant-only strategy is the best. This metric is useful to determine whether optimal rotation strategies can provide significantly better resistance management.

The second metric, resistance durability, is defined as the number of years during which the resistant-only strategy can be used without losing more than 1% of 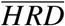, compared to the first year the resistance is used. This metric is close to the “the usefulness time” defined by Van den Bosch & Gilligan (2008), *i.e.* the number of seasons until the yield drops under a preset threshold. This metric helps assess the severity of the resistance breaking problem at hand. In the following sections, we present results for optimal periodic rotation strategies computed over an intermediate 15-season time horizon, as genetic and epidemiological parameters vary.

### Influence of fitness costs

We computed the optimal periodic rotation strategies as functions of the two fitness costs on infectiveness (*w*_*β*_) and reproduction (*w*_*r*_), to explore their effects on the two metrics defined above: the relative gain brought about by optimal periodic rotations and the resistance durability. Results are displayed in Fig. 5a.

**Fig. 5.**
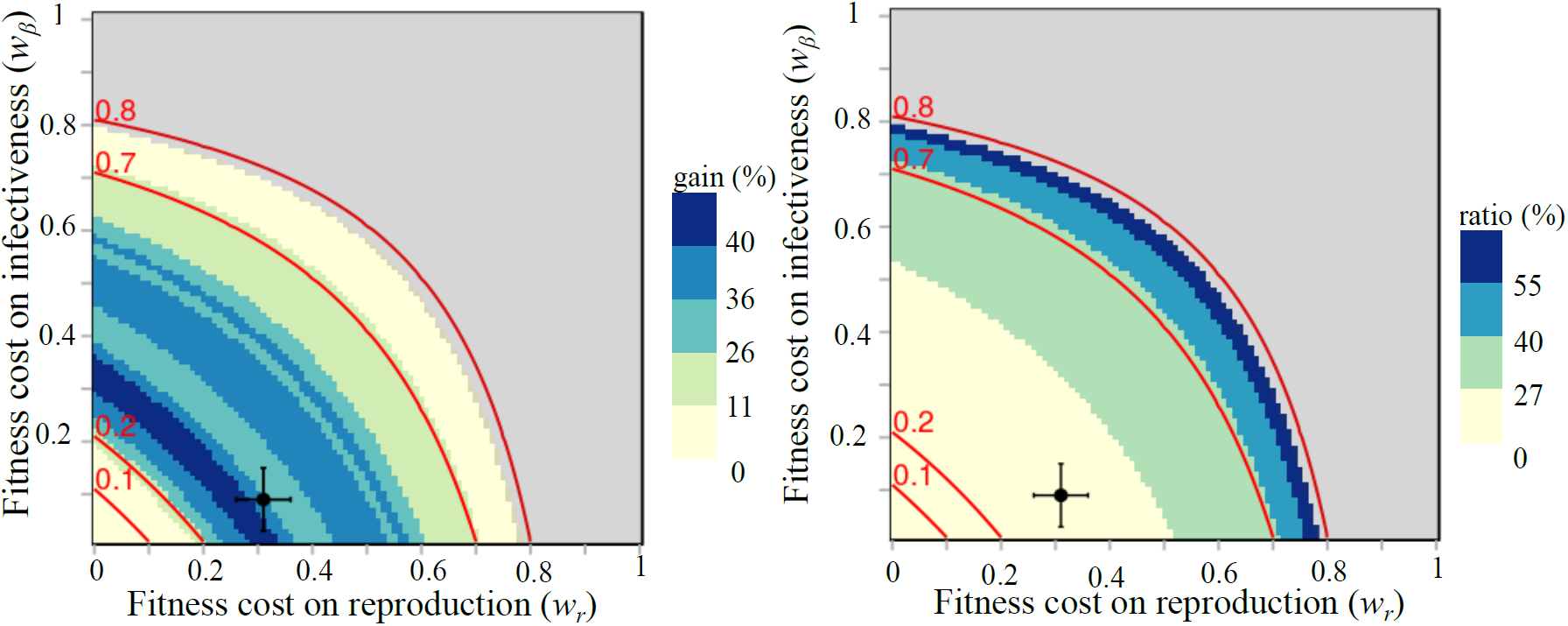
**(a)** Relative gain and **(b)** ratio of resistant plants as functions of the two fitness costs, for optimal periodic strategies computed over a 15-season time horizon. The grey area corresponds to fitness costs for which the resistance was fully durable over the 15-season time horizon. Level curves in red represent different values of the effective fitness cost *w*^*\**^ defined in (Eqn 4). The black dot and the error bars indicate the default fitness costs and their standard deviations (Castagnone-Sereno *et al.*, 2007).

The area where resistance was durable for (at least) the entire 15-season time horizon is found in the upper right part of the figure. This area corresponds to R-genes associated with very strong fitness costs of one or the other kind (*w*_*β*_ ⩾ 0.8 or *w*_*r*_ ⩾ 0.8). This means that rotation was unnecessary in such conditions, at least for the time horizon considered. For lower fitness costs, resistance was not durable and thus the use of optimal periodic rotation strategies produced a better crop yield than the resistant-only strategy (positive relative gain).

The relative gain was fairly high, except in two cases. On the one hand, when resistance breaking entailed very low fitness costs (*w*_*β*_ or *w*_*r*_ ⩽ 0.12), the relative gain was almost zero. This is not surprising since for such low fitness costs, virulent nematodes cannot be prevented from overturning the nematode population, even with rotation strategies, as they develop quite well on both resistant and susceptible plants. Cropping resistant plants is then useless and does not provide any increase in yield. On the other hand, R-genes associated with high fitness costs (*w*_*β*_ or *w* _*r*_ ⩾ 0.7) provided a relative gain of less than 10%. For such fitness costs, resistance durability was in fact quite high (12 to 14 seasons). Therefore, the resistant-only strategy was quite efficient and the additional yield provided by periodic rotations is minimal.

Significant relative gains are thus observed for R-genes inducing medium fitness costs in virulent nematodes. Relative gains can in this case reach values up to 50%. Interestingly, in the literature the fitness cost on reproduction *w*_*r*_ is estimated between 0.26 and 0.36 and the fitness cost on infectiveness *w* _*β*_ between 0.03 and 0.15, for the susceptible Saint Pierre tomato cultivar (Castagnone-Sereno *et al.*, 2007). For such realistic fitness cost values, the expected relative gain that could be realised by switching from a resistant-only strategy to an optimal periodic rotation would be between 26% and 43% with a relative gain equal to 28% for the default values parameter values.

The ratio of resistant plants deployed in the optimal periodic rotation strategies in order to achieve such relative gain values were remarkably low, lying between 13% and 27% (Fig. 5b). For the default parameter values, the ratio of resistant plants was 20%. The ratio of resistant plants used in the optimal rotation strategies increased with the values of the fitness costs.

Interestingly, Fig. 5 shows that the fitness cost distribution between infectiveness and reproduction is important for crop yield. Indeed, even though the two fitness costs had perfectly symmetrical effects, the level curves of both the relative gain and the ratio of resistant plants were markedly concave. Therefore, a balanced distribution of fitness costs (*e.g. w*_*β*_ = *w*_*r*_ = 0.4) could lead to a situation where resistance was not durable, while an uneven distribution (*e.g. w*_*r*_ = 0.8, *w*_*β*_ = 0) could lead to a durable situation. The two fitness costs thus did not act in an additive manner and interacted negatively. The derivation of the multiseason basic reproduction number *R*_0_ of virulent nematodes revealed that it depended only on the product (1 − *w*_*β*_)(1 − *w*_*r*_) (Supporting Information Methods S1). We hence defined an “effective” fitness cost as:

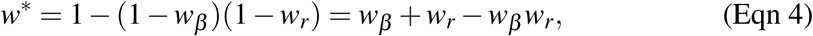

whose level curves perfectly reflected the level curves of the relative gain and ratio of resistant plants (Fig. 5). The performance of resistance-based strategies therefore appeared to be entirely determined by this quantity.

In the following, we thus present results in terms of this effective fitness cost *w*^*\**^.

### Interplay between epidemiological scenarios and genetic parameters

We studied the influence of the genetic parameters in interaction with the epidemiological scenarios on the relative gain and durability. Fig. 6 shows the relative gain obtained for a 15-season time horizon as a function of the effective fitness cost (*w*^*\**^), for different values of the fraction of virulent offspring (*δ*) and the four epidemiological scenarios. Parameter ranges ensuring resistance durability over the 15-season time horizon were identified (grey areas). *δ* had no effect on durability according to our definition. Indeed, when only resistant plants were deployed, avirulent nematodes could not reproduce. The resistance was durable as the effective fitness cost *w*^*\**^ overshooted a given threshold, which strongly increased with the severity of the epidemic scenario. For instance, for low aggressiveness of the nematode, R-genes associated with effective fitness costs between 0.20 and 1 were durable (Fig. 6a), while in the extreme scenario, they were durable only for fitness costs larger than 0.95 (Fig. 6d).

**Fig. 6.**
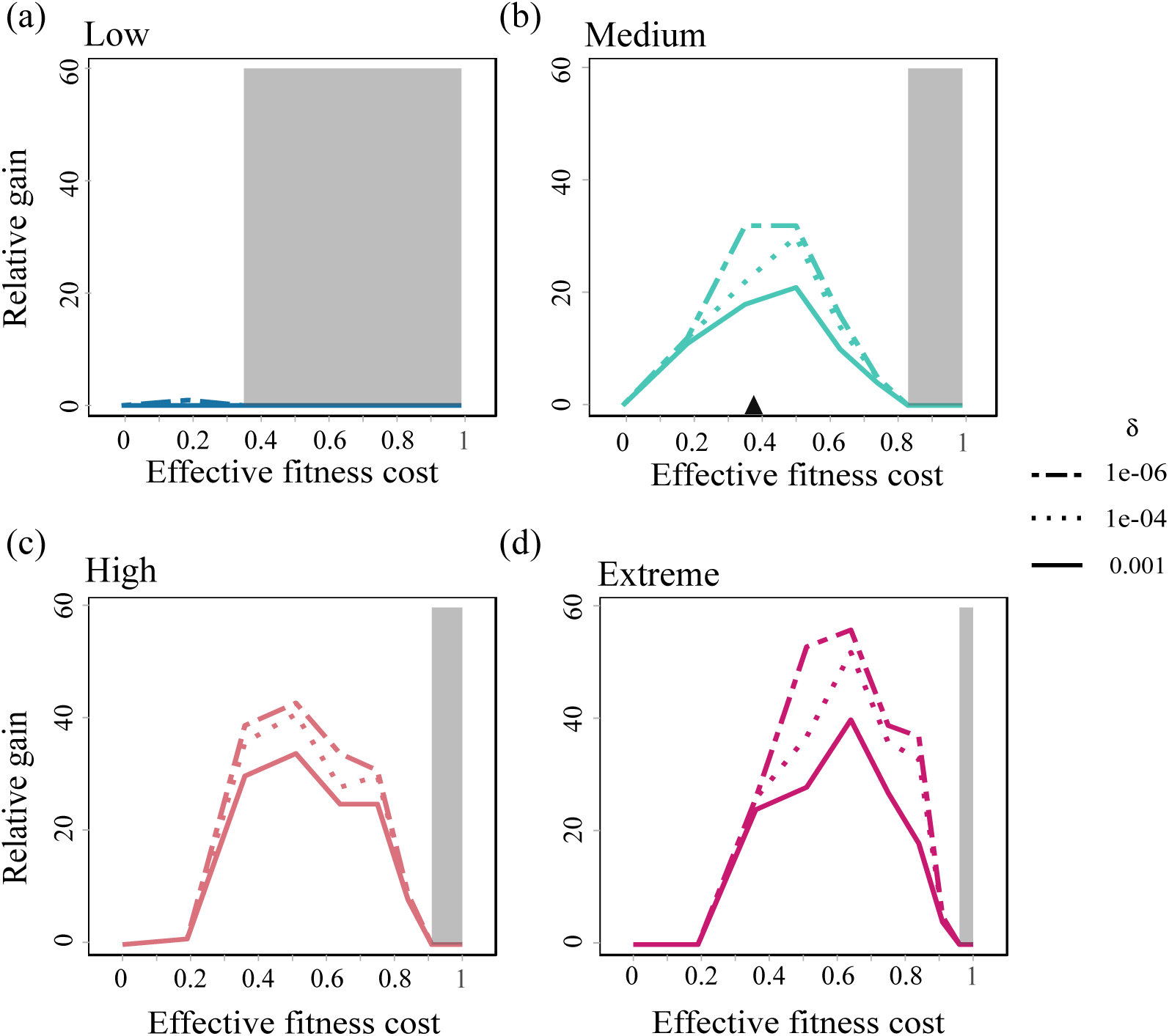
Graphical representation of the relative gain for a 15-season time horizon for the four epidemiological scenarios (a-d) defined in Table 2, as a function of the effective fitness cost (*w*^*\**^) and the fraction of virulent offspring (*δ*). The default effective fitness cost *w*^*\**^ = 0.37 is represented by the black triangle (Castagnone-Sereno *et al.*, 2007). Grey areas represent the values of *w*^*\**^ for which the resistance was durable over the 15-season time horizon.

The relative gain varied significantly according to the genetic parameters and epidemiological scenarios, except for the low epidemiological scenario where it remained close to zero (Fig. 6a). In this case, nematode infestation remained very low so that the resistant-only strategy actually provided very good control. The relative gain increased with nematode aggressiveness and decreased with the fraction of virulent offspring *δ*. The best gains were found for R-genes associated with medium to high effective fitness costs(between 0.4 to 0.65). For example, an extreme nematode aggressiveness combined with a low fraction of virulent offspring *δ* = 10^−6^ and a fitness cost *w*^*\**^ = 0.65 yielded a relative gain of up to 58% (Fig. 6d). Hence, nematode aggressiveness tended to increase the advantages of cultivar rotations over the resistant-only strategy.

### Robustness of deployment strategies

Finally, we tested the robustness of the optimal periodic rotation strategies by confronting their efficacy to variations in parameter values. Fig. 7 represents the relative gain when applying the optimal periodic strategy computed over a 15-season time horizon for three epidemiological scenarios, in response to 10% parameter variations.

**Fig. 7.**
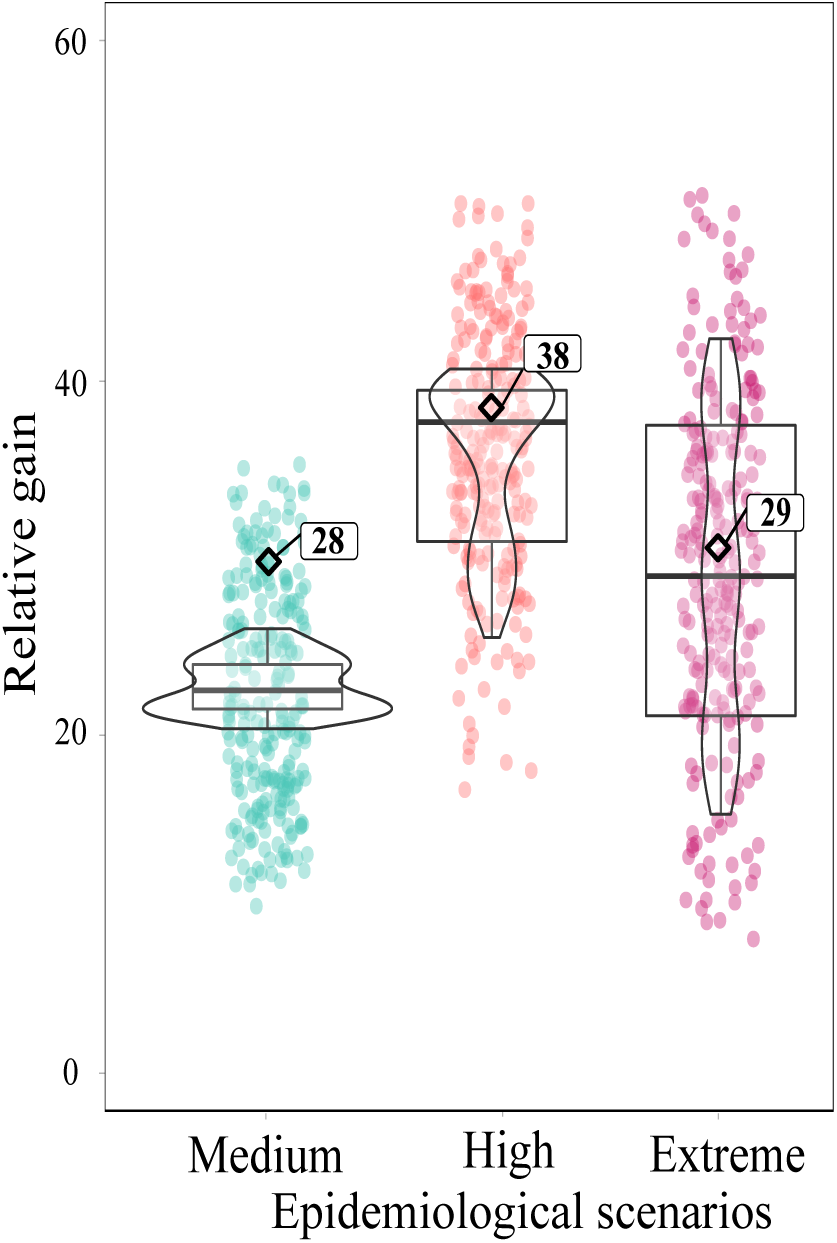
Robustness of the relative gain to changes in model parameters for three epidemiological scenarios: medium, high, extreme. For each scenario, the optimal periodic strategy computed over a 15-season time horizon for the default parameters values (Tables 1–2) was applied. The relative gain obtained with the default parameter values (black diamond) and 243 combinations derived from *±*10% variations in key parameters (coloured dots) are plotted, as well as the corresponding boxplots.

The relative gain remained positive, but was globally sensitive to the parameter variations tested. For the medium epidemiological scenario, most parameter combinations yielded a relative gain lower than the 28% gain obtained with the default parameter values (black diamond in Fig. 7). For the other scenarios, combinations yielded lower and higher relative gains.

However, for all epidemiological scenarios and all parameter combinations, the optimal periodic strategy remained more efficient than the resistant-only strategy, since the relative gain remained positive. For instance, the 1*R* + 5*S* strategy always performed better than a resistant-only strategy over a 15-season time horizon for the medium epidemiological scenario. In that sense, the relative performance of the optimal periodic strategy was globally very robust to parameter changes.

## Discussion

### Crop rotation is an efficient strategy

The present study was based on a new model of plant–nematode interactions parameterized from the literature and fitted to experimental data, so as to be representative of the tomato–RKN system. As a key result, we found that alternating susceptible and resistant plant cultivars in time can help limit the proportion of virulent nematodes in RKN populations and thereby reduce crop loss importantly. According to our simulations, relative gains as high as 40% can be achieved, compared to the baseline strategy of deploying only resistant plants, over time horizons of 15 years or more.

The relative gain achievable with optimal crop rotations was found to be greatest in high or extreme epidemiological scenarios, i.e. for very aggressive nematodes. The latter result echoes previous findings on the influence of epidemic intensity on resistance durability in the context of spatial mixtures (Van den Bosch & Gilligan, 2003; Fabre *et al.*, 2012). The gain also increased, to a smaller extent, if the fraction of virulent offspring in avirulent egg-clutches is smaller, and if the culture is sustained over longer temporal horizons. Remarkably, the relative gain obtained from virulence costs similar to those estimated for the *Mi-1* resistant gene is close to the maximum achievable gain value Fig. 5a, suggesting that crop rotation is a particularly promising strategy when deploying *Mi-1* cultivars. We also found that periodic crop rotation strategies are almost as effective as free (unconstrained) alternation strategies. This result has considerable importance, since periodic rotation patterns are in real-world applications much easier for crop growers to implement than complicated unconstrained sequences.

Few recent theoretical studies have considered the deployment of different cultivars over time (alternation strategies). One is Rimbaud *et al.* (2018), that compared four resistance deployment strategies of major resistance genes: mosaics, mixtures, rotations and pyramiding, to manage cereal rust fungi in agricultural landscapes durably. They found cultivar rotation to be the most efficient in the long-term, once every R-genes had been overcome. In a study of plant virus epidemic control by mixing resistant and susceptible cultivars in space and time, Fabre *et al.* (2015) identified that in more than 20% of the scenarios considered, optimal strategies involved cultivar rotation at the landscape scale. Studies are even scarcer regarding RKN, for which the literature on cultivar rotation is essentially experimental. For these low-dispersing soil-borne pests, data support our modelling predictions in suggesting that rotations are an effective way to reduce yield losses and to delay outbreaks (Tzortzakakis *et al.*, 2000; Miller *et al.*, 2006; McSorley, 2011). For instance, Djian-Caporalino *et al.* (2014) experimentally compared the performance of several strategies to control RKN in vegetable cropping systems, including rotations of two major R-genes in pepper cultivars, over three years. They reported that cultivar rotation can improve epidemiological control and resistance durability. Another study by Talavera *et al.* (2009) on RKN management compared the effects of four crop rotations between resistant and susceptible tomato plants in a three-year field experiment. Regarding crop yield and durability, this study showed that the best strategy consisted in growing two resistant cultivars, followed by one susceptible cultivar. This is strikingly consistent with our modelling predictions, since we found that the yield-maximising strategy, over a three-season temporal horizon, is 1*R* + 2*S* Fig. 4b. Our modelling results further indicate that the performance of crop rotations for RKN control would be even more pronounced over longer time horizons.

### Crop rotation (usually) requires low ratios of resistant plants

Interestingly, the optimal rotation strategies identified in this study were characterised by relatively low ratios of resistant plants, as soon as the temporal horizon exceeded seven cropping seasons (Fig. 5b). Since avirulent nematodes thrive on susceptible plants, low ratios of resistant plants are expected to increase crop loss, especially in the short-term. However, in the longer term, low ratios limit selection for virulent variants, thus prolongating the efficacy of resistant plants when those are deployed. For RKNs, it appears that the relatively fast within-season epidemiological dynamics sets the optimal balance between the two effects at a low ratio of resistant plants. Our results are consistent with Van den Bosch & Gilligan (2003), who showed that, in many instances, low ratios allowed to make the most of resistance, by reducing the selection pressure for virulent pathogens and promoting resistance durability.

Interestingly, studies of spatial deployment strategies tend to report higher optimal ratios of resistant plants. Fabre *et al.* (2012), working on plant resistance to viruses, demonstrated that optimal ratios were frequently over 50%. For instance, for low fitness costs, the ratio ranged between 50 and 70%, depending on the epidemic profile. Regarding phytopathogenic fungi, Papaïx *et al*. (2014) also found that high ratios combined with low levels of variety aggregation provided optimal control of the fungi in agricultural landscapes. Therefore, the selection pressure in favour of virulent variants seems to be lower when mixing resistant and susceptible cultivars in space compared to alternating them over time.

It should be remarked that low ratios of resistant plants are in total contrast with the currently dominant agricultural practices, based on the regular cropping of tomato cultivars bearing the same Mi-1 resistance gene. Be it in the field or in experimental studies, such resistant-only strategies often fail, and virulent RKNs overcoming resistance have been observed in most tomato growing areas worldwide (Seid *et al.*, 2015). More specifically, experimental findings have shown that three consecutive cropping seasons of the *Mi-1* gene in tomatoes were enough for RKNs to overcome the resistance (Eddaoudi *et al.*, 1997; Verdejo-Lucas *et al.*, 2009). These findings are consistent with our results when fitness costs are not too severe, close to available experimental estimates (Castagnone-Sereno *et al.*, 2007; Djian-Caporalino *et al.*, 2011). The intense deployment of resistant cultivars is thus bound to cause boom and bust cycles in this system (Brown & Tellier, 2011). During the boom, crop yield increases rapidly thanks to the use of new resistant cultivar by growers and farmers. Nevertheless, it is followed by a bust, characterised by the rapid breakdown of the resistance by virulent variants and a drop in crop yield. The switch to a new cultivar, carrying a fresh resistance gene, then triggers a new cycle. To break this cycle and preserve the efficiency of resistance genes, which are scarce and valuable resources, cultivar rotations such as the ones proposed in this study are a feasible and sustainable alternative.

### What makes a good resistance gene?

We investigated the effects of varying three mechanistic parameters characterizing how a resistance gene behaves with respect to resistance breaking by RKNs: the fitness cost it imposes on the infectivity of virulent nematodes (*w*_*β*_), the fitness cost it imposes on their reproduction (*w*_*r*_), and the frequency of virulence appearance in avirulent clutches (*δ*). Obviously, one would seek R-genes that, when overcome, would generate high values of the first two parameters, and low values of the third.

Our results showed that the two fitness costs had interchangeable effects in shaping the population dynamics of virulent variants. However, the two costs interacted negatively, as the benefit of increasing one fitness cost was reduced when the other fitness cost is already high (Fig. 5a). This original result entails that, when evaluating the potential of resistance genes to improve durability, breeders should seek and introgress R-genes with maximal fitness cost on either one or the two components of the nematode life-cycle (reproduction or infectivity), rather than a balanced distribution of the two types of costs. To help address the existence of two different types of fitness costs, a specificity of our model, we derived a simple formula to synthesize the two fitness costs into one effective fitness cost, according to which different resistance genes can be ranked in terms of their durability. Comparatively, the rate of production of virulent nematodes *δ* had virtually no impact on the durability of resistance genes.

Rotation strategies provided the largest relative benefits over the resistant-only strategy for intermediate fitness costs. Such measurements are not always easily accessible in the literature, but this property seems to hold in a few other studies. For instance, a reinvestigation of the simulation data on plant virus epidemics obtained by Fabre *et al.* (2012) for high epidemic intensities showed that the best relative gains were obtained for intermediate fitness costs. Another study by Rousseau *et al.* (2019) showed that relative additional gains, provided by combining quantitative and qualitative resistances over qualitative resistances only, were most noticeable for intermediate fitness costs. In both studies, the reasons for this were similar to the present study: high fitness costs induced durable resistance so that the yield could only be marginally increased, whereas low costs induced poorly efficient resistance that did not benefit from an optimal deployement. R-genes associated with intermediate fitness costs are thus the ones that could benefit the most from improvements in terms of deployment or cultivar genetic background.

### Optimal rotations in practice

A major outcome of this work would be to recommend custom optimal resistance deployment strategies to crop growers, depending on the temporal horizon sought, but also on the epidemic context, the R-genes to be deployed, and on the agricultural practices that determine model parameter values. However, soil infestation, and epidemiological or genetic parameters are particularly difficult to estimate, and likely subject to considerable uncertainty. For instance, Djian-Caporalino *et al.* (2011) found a large variability in the fitness costs on reproduction. To address this issue, we simulated the use of optimal periodic strategies, as computed for reference parameters values, and investigated how their performance responded to parameter variations. We found that the relative gain was globally robust to parameter changes. Thus, optimal periodic rotations can outperform the resistant-only strategy in terms of crop yield even if the relevant parameters are known imperfectly, which is a very desirable property in practice. There are still few studies that investigate the robustness of resistance deployment strategies, or more generally plant pathogen control methods. A similar analysis to parameter misspecification was conducted by Hyatt-Twynam *et al.* (2017) to assess the performance of optimal strategies to control the spread of citrus canker in Florida, using one at a time epidemiological parameter changes. In the context of fungicide resistance management, Elderfield *et al.* (2018) found that mixtures always outperformed alternations when parameters varied, but not the deployment strategy. More such studies should arise to help bridge the gap between theoretical resistance deployment strategies and their implementation in the field. Optimal strategies could feature high year-to-year variations in yield, which may not be economically viable for farmers. Taking advantage of the limited mobility of nematodes, this issue could be adressed by implementing asynchronous crop rotation strategies in different rows or plots, provided that contamination between those be carefully avoided. The seasonal yield variations in each row would average out, ensuring a more stable income for farmers while achieving the performance of the optimal rotation strategy. Currently, *Mi-1* is the only resistance gene used in tomato cultivars, which makes our study entirely relevant for the tomato-RKN pathosystem. However, our model could readily be extended to incorporate different R-genes and serve as a basis to evaluate more complex resistance deployment strategies, involving rotations between susceptible and several resistant cultivars, including pyramided ones.

## Acknowledgements

This work was funded by INRA and Région Sud PACA, France. It was certified by the Terralia competitiveness research cluster. The authors are grateful to the Inria Sophia Antipolis – Méditerranée “NEF” computation cluster for providing resources and support.

## Author Contribution

VC, LM and ST conceived the research, with input from PC-S and CD-C. SN conducted the research, with input from all co-authors. MM and TP contributed to the research. SN performed the analysis, with input from VC, LM and ST. SN wrote the manuscript, with input from VC, LM, ST. The manuscript was reviewed by PC-S and CD-C.

## Supporting Information

**Fig. S1** Diagram representing the plant-nematode interaction model for two successive cropping seasons of resistant and susceptible plants.

**Fig. S2 (within Methods S3)** Global sensitivity indices on the healthy root density (a yield proxy) for a susceptible-only strategy over a 15-season time horizon.

**Methods S1** Computation of the season to season basic reproduction numbers *R*_0_ for avirulent and virulent nematodes.

**Methods S2** Model fitting to experimental data describing the infection dynamics of susceptible tomato roots by avirulent nematodes during a cropping season.

**Methods S3** Sensitivity analysis to assess the parameter impact on the healthy root density (a yield proxy) for a susceptible-only strategy over a 15-season time horizon.

