## Supporting Information for "A multi-seasonal model of plant-nematode interactions and its use to identify optimal plant resistance deployment strategies"

The following Supporting Information is available for this article:

**Fig. S1** Diagram representing the plant-nematode interaction model for two successive cropping seasons of resistant and susceptible plants.

**Fig. S2 (within Methods S3)** Global sensitivity indices on the healthy root density (a yield proxy) for a susceptible-only strategy over a 15-season time horizon.

**Methods S1** Computation of the season to season basic reproduction numbers $R_0$ for avirulent and virulent nematodes.

**Methods S2** Model fitting to experimental data describing the infection dynamics of susceptible tomato roots by avirulent nematodes during a cropping season.

**Methods S3** Sensitivity analysis to assess the parameter impact on the healthy root density (a yield proxy) for a susceptible-only strategy over a 15-season time horizon.

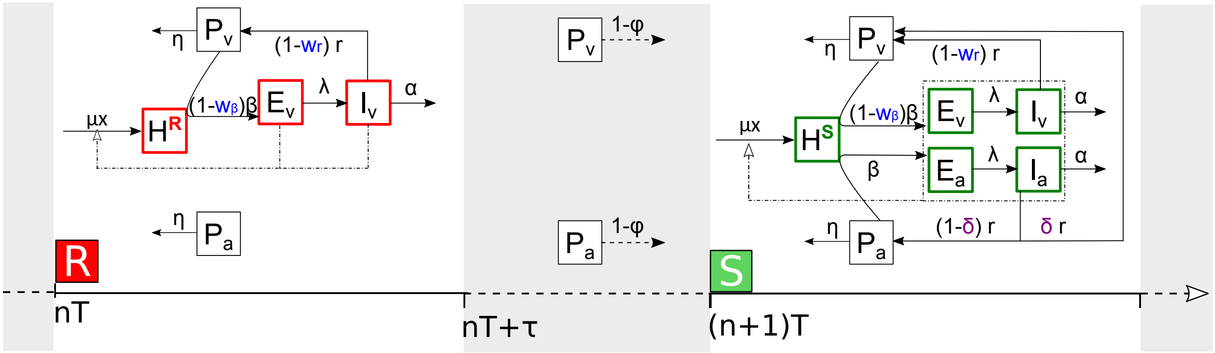


**Fig. S1** Compartmental diagram representing the plant-nematode interaction model for two successive cropping seasons of resistant (superscript $X = R$) and susceptible (superscript $X = S$) plants. Healthy plant roots ($H^X$) are infected by virulent (subscript $v$) and avirulent (subscript $a$) nematodes in the soil ($P$), before becoming latently infected ($E = E_a + E_v$) and then infectious ($I = I_a + I_v$) feeding sites. Between cropping seasons (shaded area), free living nematodes remain in the soil. Parameters are described in Table 1 (main text).

### Methods S1. Computation of the season to season basic reproduction numbers $R_0$

Season to season basic reproduction numbers quantify the multiplication rate of a pathogen from the beginning of a cropping season to the beginning of the next season, when pathogen densities are vanishingly low (Mailleret *et al.*, 2012).

To evaluate the basic reproduction number of avirulent nematodes $P_a$ on susceptible plants, we assumed that the dynamics of infected feeding sites $E_a$ and $I_a$ were fast and that the fraction of virulent offspring $\delta$ tended to 0. Under these approximations, the dynamics of free living nematodes $P_a$ becomes:

$$\dot{P}_a(t) = P_a(t)\left(\beta\left(\frac{\varepsilon_a^S r}{\alpha} - 1\right)H^S(t) - \eta\right),$$

so that:

$$P_a(\tau) = \exp\left(\int_{t=0}^{t=\tau}\left(\beta\left(\frac{\varepsilon_a^S r}{\alpha} - 1\right)H^S(t) - \eta\right)dt\right)P_a(0).$$

Healthy roots grow linearly in the absence of nematodes so that at time $t$ during the course of a season $H^S(t) = H_0 + \mu x t$. Therefore:

$$P_a(\tau) = \exp\left(\beta\left(\frac{\varepsilon_a^S r}{\alpha} - 1\right)\left(H_0\tau + \frac{\mu x \tau^2}{2}\right) - \eta\tau\right)P_a(0).$$

Taking into account between-season survival $\varphi$, the season to season basic reproduction number of avirulent nematodes on susceptible plants therefore reads:

$$R_{0,a}^S = \varphi \exp\left(\beta\left(\frac{\varepsilon_a^S r}{\alpha} - 1\right)\left(H_0\tau + \frac{\mu x \tau^2}{2}\right) - \eta\tau\right).$$

One can proceed in a very same way to compute the basic reproduction number of avirulent nematodes on resistant plants, accounting for the absence of free living nematodes produced during such a season. This yields:

$$R_{0,a}^R = \varphi \exp\left(-\beta\left(H_0\tau + \frac{\mu x \tau^2}{2}\right) - \eta\tau\right),$$

which is clearly lower than 1.

Finally, since virulent nematodes develop similarly on resistant and susceptible plants, their basic reproduction number is the same on both plants. Applying the method described above and taking into account the presence of fitness costs $w_\beta$ and $w_r$, this leads to:

$$R_{0,v} = \varphi \exp\left(\beta\left(\frac{(1-w_\beta)(1-w_r)\varepsilon_v r}{\alpha} - 1\right)\left(H_0\tau + \frac{\mu x \tau^2}{2}\right) - \eta\tau\right). \qquad \text{(Eqn S1)}$$

Introducing the effective fitness cost $w^* = 1-(1-w_\beta)(1-w_r)$, one gets:

$$R_{0,v} = \varphi \exp\left(\beta\left(\frac{(1-w^*)\varepsilon_v r}{\alpha} - 1\right)\left(H_0\tau + \frac{\mu x \tau^2}{2}\right) - \eta\tau\right).$$

**Mailleret L**, **Castel M**, **Montarry J**, **Hamelin FM**. **2012**. From elaborate to compact seasonal plant epidemic models and back: Is competitive exclusion in the details? *Theoretical Ecology* **5**: 311–324.

### Methods S2. Model fitting to experimental data

Three parameters could not be set from published data: the infection rate ($\beta$), the conversion factor between root biomass and density of feeding sites ($x$) and the plant growth scaling factor ($k$). To estimate these parameters, data obtained by Ehwaeti *et al.* (1998) were used. They describe the infection dynamics of susceptible tomato (cv *Moneymarker*) roots by avirulent nematodes *Meloidogyne incognita*, which were initially inoculated in the soil at five controlled densities ($P_0(i)$ with $i = 1,\ldots,5$). The nematode density in the roots ($N_{\text{data}}$) was measured after 42 days and 135 days of cultivation. The relative root biomass ($B_{\text{data}}$), *i.e.* the root biomass of an infected plant divided by the root biomass of an uninfected plant, was also measured. Both measures were reported as a mean over five replicates, for each initial nematode density. Only final measures after 135 days of cultivation were used to estimate the parameters (measures after 42 days were used to assess the model validity).

Model fitting was performed by finding the $x,k,\beta$ and $y$ parameter values that minimised the distance between the model output and the data using on a weighted square metric:

$$J(x,k,\beta) = \sum_{i=1}^{5} \left( \frac{N_{\text{model}}(i,x,k,\beta) - N_{\text{data}}(i)}{\text{mean}_i(N_{\text{data}}(i))} \right)^2 + \sum_{i=1}^{5} \left( \frac{B_{\text{model}}(i,x,k,\beta,y) - B_{\text{data}}(i)}{\text{mean}_i(B_{\text{data}}(i))} \right)^2.$$

Parameter $y$ was introduced to take into account the extra root mass corresponding to the root-knot galls caused by nematodes. It was estimated along with the three other parameters. $N_{\text{model}}(i,x,k,\beta)$ and $B_{\text{model}}(i,x,k,\beta)$ were computed, based on model (Eqn 1, main text) integrated at time $t = 135$ days, as follows:

$$\begin{aligned} N_{\text{model}}(i,x,k,\beta) &= E_a(135) + I_a(135), \\ B_{\text{model}}(i,x,k,\beta,y) &= \frac{H^S(135) + yE_a(135) + yI_a(135)}{H^{S*}(135)}, \end{aligned}$$

with the initial nematode density in the soil $P_a(0) = P_0(i)$, except for $H^{S*}$ which was computed with $P_a(0) = 0$, and with given parameters $x,k,\beta$ and $y$ (remaining parameter values can be found in Table 1, main text).

Division by the means made both $N$ and $B$ terms dimensionless so they could be summed. The computation of the parameter values that minimised $J$ was achieved using the optim function of R with the default Nelder-Mead method.

**Ehwaeti ME**, **Phillips MS**, **Trudgill DL**. **1998**. Dynamics of damage to tomato by *Meloidogyne incognita*. *Fundamental and Applied Nematology* **21**: 627–635.

### Methods S3. Sensitivity analysis

We performed a global sensitivity analysis to assess the parameter impact on the healthy root density ($\overline{HRD}$) for a susceptible-only strategy over a 15-season time horizon. The method used is based on factorial design and analysis of variance (ANOVA). The $\overline{HRD}$ was considered as the output (or observation), the parameters as factors.

We first explored the parameter space by varying 10 epidemiological parameters ($P_0$, $p_v$, $\beta$, $w_\beta$, $r$, $w_r$, $\delta$, $\alpha$, $\eta$, $\varphi$) with three levels per parameter: the reference value found in the literature or estimated according to Supporting Information Methods S2 and $\pm 30\%$ variations, except for $P_0$ for which larger variations were chosen (Table 1, main text). A full factorial design, defined as all possible combinations of the three parameter levels, corresponds to $3^{10} = 59049$ combinations and would have required the same number of simulations to compute the corresponding $\overline{HRD}$. To reduce this number, we implemented a fractional factorial design (a subset of the full design), chosen in order to estimate all parameter main effects and two-way interactions. The fractional factorial design was obtained using the PLANOR R package (https://CRAN.R-project.org/package=planor) and consisted of 243 parameter combinations, yielding as many simulations.

By means of an ANOVA, we then proceeded with the observed variance decomposition and estimated the sum of squares associated with each factorial term, main effect $SS_i$ or the two-way interaction $SS_{i,j}$. According to the sparsity-of-effects principle, a system is usually dominated by main effects and low order interactions, so neglecting higher interactions can still provide good estimates. Denoting by $SS_T$ the total sum of squares, the total sensitivity index of parameter $p_i$ is defined as follows:

$$tSI_i = \frac{\overbrace{SS_i}^{\text{main effect}} + \overbrace{\sum_{j \neq i} SS_{i,j}}^{\text{two-way interactions}}}{SS_T}. \qquad \text{(Eqn S2)}$$

It represents the fraction of the output variability explained by parameter $p_f$. We used the MULTISENSI R package (https://CRAN.R-project.org/package=multisensi) for this analysis.

The results of the global sensitivity analysis are shown in Fig. S2. Interactions had little impact, so parameter main effects largely explained the output variability. Parameters that most influenced the $\overline{HRD}$ were the infection rate $\beta$ ($tSI_\beta = 30\%$ ), the nematode reproduction rate $r$ ($tSI_r = 28\%$), as well as the nematode mortality rates in the soil $\eta$ ($tSI_\eta = 23\%$) and in the plant $\alpha$ ($tSI_\alpha = 18\%$). The other parameters had little impact on the output.

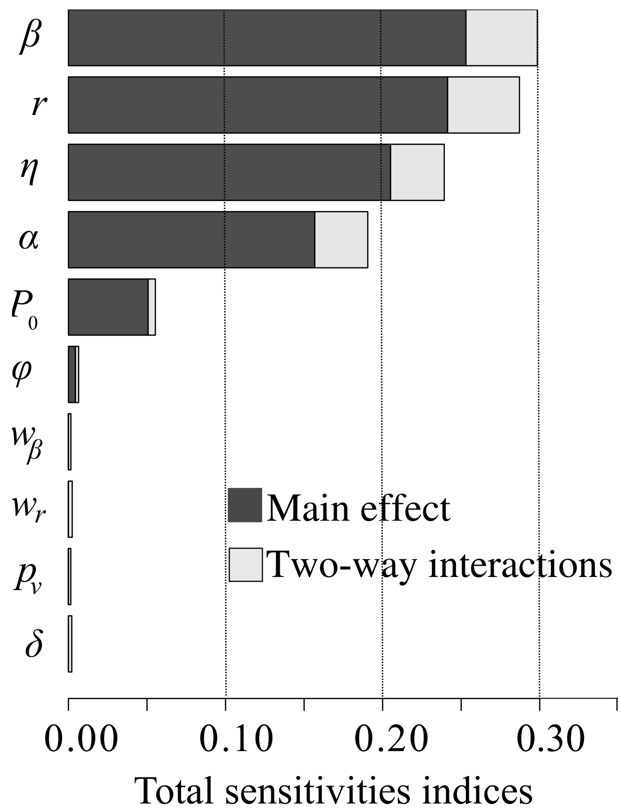


**Fig. S2** Global sensitivity indices on the healthy root density ($\overline{HRD}$) for a susceptible-only strategy over a 15-season time horizon. Total sensitivity indices for the 10 epidemiological parameters are ranked in descending order and split in main effect (black bar) and two-way interactions (grey bar) according to (Eqn S2).
